# Genetic control of cellular morphogenesis in Müller glia

**DOI:** 10.1101/392902

**Authors:** Mark Charlton-Perkins, Alexandra D. Almeida, Ryan B. MacDonald, William A. Harris

## Abstract

Of all the cells in the body, those with the greatest variety of shapes reside in the central nervous system yet they all start their postmitotic lives as simple elongated cells of the neuroepithelium^1^.The molecular processes by which these, or indeed any, cells gain their particular cell-specific anatomies remain largely unexplored.We, therefore, developed a strategy to identify the genes involved in cellular morphogenesis using Müller glial (MG) cells in the vertebrate retina as a model system.These radially oriented cells, discovered by Heinrich Müller in 1851 and named in his honour^2^, are astonishingly complex yet, as the great neurohistologist Ramon y Cajal first noted, they share a conserved set of key anatomical features^3^.Using genomic and CRISPR based strategies in zebrafish, combined with a temporal dissection of the process, we found more than 40 genes involved in MG cell morphogenesis.Strikingly, the sequential steps of anatomical feature addition are regulated by successive expression of cohorts of interrelated genes, revealing unprecedented insights into the developmental genetics of cellular morphogenesis.

---

Despite their species-specific variability, mature MG cells in all vertebrate retinas share the following key features (Figure 1A): 1) their cell bodies sit in the middle cellular layer of the retina - the inner nuclear layer (INL); 2) their central radial stalks span the apicobasal extent of the retina, with endfeet upon both the outer and the inner limiting membranes (OLM and ILM); 3) fine branches emerge laterally from these central stalks extending differentially into the two synaptic layers, known as the outer and inner plexiform layers (OPL and IPL).Thus, the mature MG cell morphology facilitates their contact with every cell, and possibly every synapse, in the retina and enables them to carry out their many homeostatic physiological functions^4,5^.

**Figure 1.**
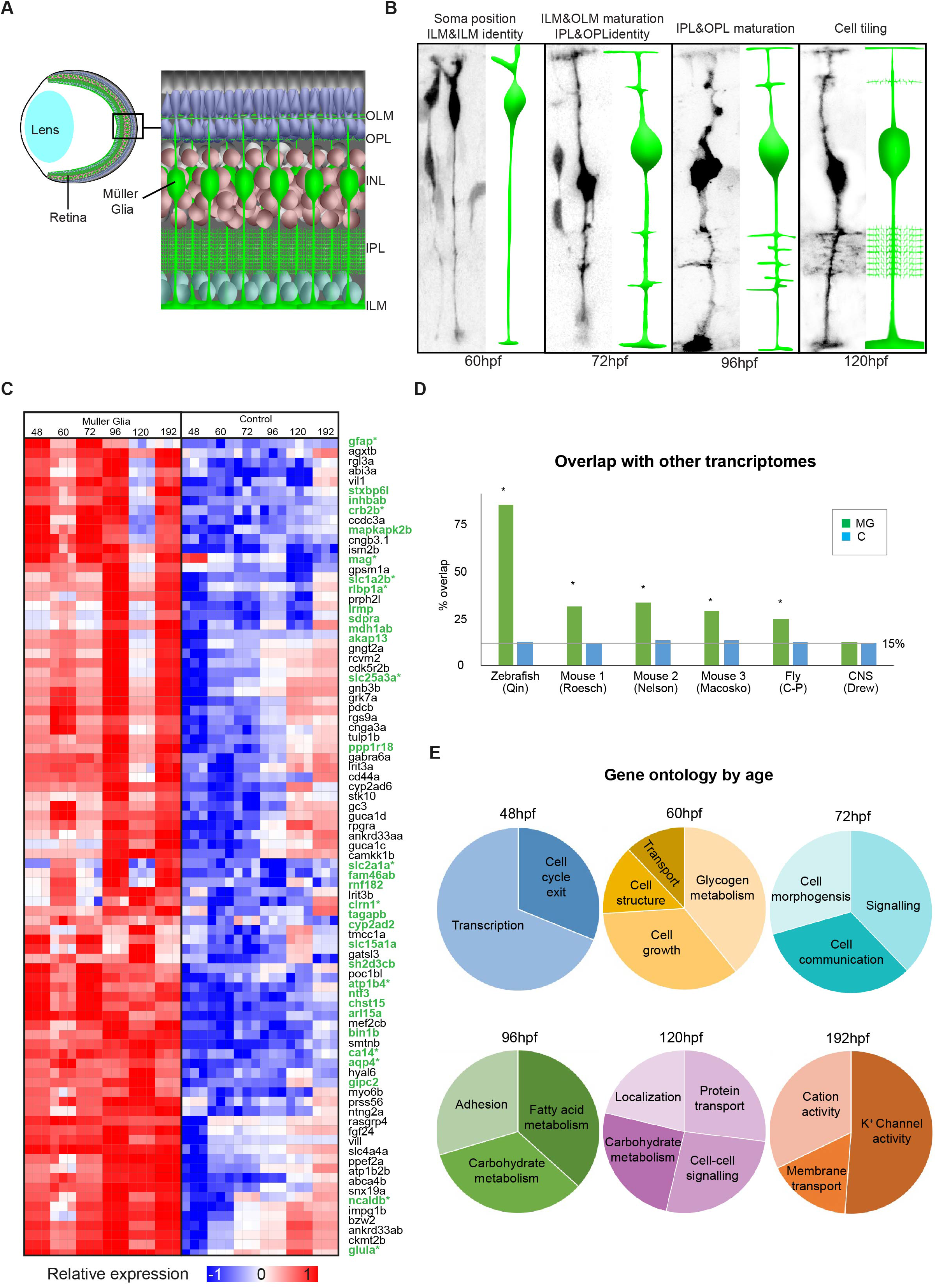
Temporal MG cell morphology and gene expression. A)Diagrammatic representation of the retina within the eye showing positioning of MG cells.B) Tg(TP1:Venus) transplanted MG cells showing the time course of MG cell differentiation that gives rise to the distinct MG compartments (OLM – outer limiting membrane, OPL – outer plexiform layer, INL – inner nuclear layer, IPL – inner plexiform layer, ILM – inner limiting membrane).C) Heatmap of top 100 significantly expressed genes with glial genes in green (* indicates previous reported expression in MG).D) Overlap of zebrafish MG enriched genes with previously reported MG transcriptomes from zebrafish, mouse and fly ^24,25,27,54,55^. * -indicates significance (Bonferroni adjusted p-value <0.001) by Fisher’s exact test.E) Representative gene ontology proportions of MG genes enriched at 48, 60, 72, 96 and 120hpf.

The conserved, yet complex cellular anatomy of MG cells makes them an excellent cell type to investigate the genes involved in cellular morphogenesis.Previous studies have shown that MG exhibit several discrete steps of anatomical specialization^3,6–11^.Notch signalling is essential for MG cell specification and in the *Tg(TP1:Venus)* transgenic line^12^, the Notch-responsive element TP1 drives expression of the fluorescent protein Venus, allowing MG cells to be followed from the time of their initial specification in zebrafish at ~60 hours post-fertilisation (hpf)^7^.We visualised MG morphogenesis in zebrafish *in vivo,* by transplanting blastomeres from the transgenic zebrafish line, into wild-type hosts (Figure 1B).At 60hpf, the MG cell bodies begin to migrate basally to their stereotypic position in the middle of the INL of the retina^7^.By 72hpf, they begin to expand their apical and basal endfeet along the OLM and ILM respectively and they extend dynamic filopodia from their central stalks, which identify the OLP and the apical and basal limits of the IPL.By 96hpf they elaborate fine processes with the plexiform layers^8^.One of the last steps in this process is that MG cells space themselves out across the retina.When first specified, MG cells are positioned much more randomly but, like many types of retinal neurons, their processes are arranged in tiled mosaic with little overlap between the domains of neighbouring MG cells^8,9^.Homotypic repulsive cell interactions are thought to account for this^8,13^, as focal ablation of MG cells results in nearby MG cells extending processes to fill in the spaces previously occupied by the ablated MG cell^8^.Thus, by the time robust vision commences in zebrafish at about 120hpf^14^, MG cells have gained a full set of cell-specific anatomical characteristics (Figure 1A, B)^7,8^.

Cellular morphogenesis has been studied mostly at the cellular level, and in spite of the obvious coupling with development stages, the genetic control of cell shape in development is very poorly understood.To search for the genes involved in MG cell morphogenesis, we first identified genes expressed preferentially in FACS-sorted MG at specific times that span this morphogenetic process (48hpf, 60hpf, 72hpf, 96hpf, 120hpf and 192hpf) (Figure 1C).Hierarchical clustering and principal component analysis of these data reveal that replicates of individual time-points have no apparent differences (Extended data fig 1A).However, significant differential gene expression is notable over the course of MG cell differentiation (Extended data fig 1A; Supp table 1).Importantly, clustering all time-points by differential gene enrichment (FDR), shows that several of the top 100 genes (e.g.*gfap, slc1a2b, rlbp1a, apq4, slc25a3a, slc2a1a, ncaldb, glula* and *slc1a2a)* have previously been associated directly with MG cells or other glia (Figure 2C, Supp table 2)^15–23^.Furthermore, our data has greater than 80% overlap with a previous zebrafish MG cell transcriptome dataset (Figure 1D)^24^.The conservation of MG cell biology is highlighted by the significant genetic overlaps with three studies of mouse MGs (Fig 1D; Extended data fig 1B)^25–27^.Finally, invertebrate retinal glia also shares a significant proportion of gene orthologs with mouse MG cells (Figure 1D)^28^.

**Figure 2.**
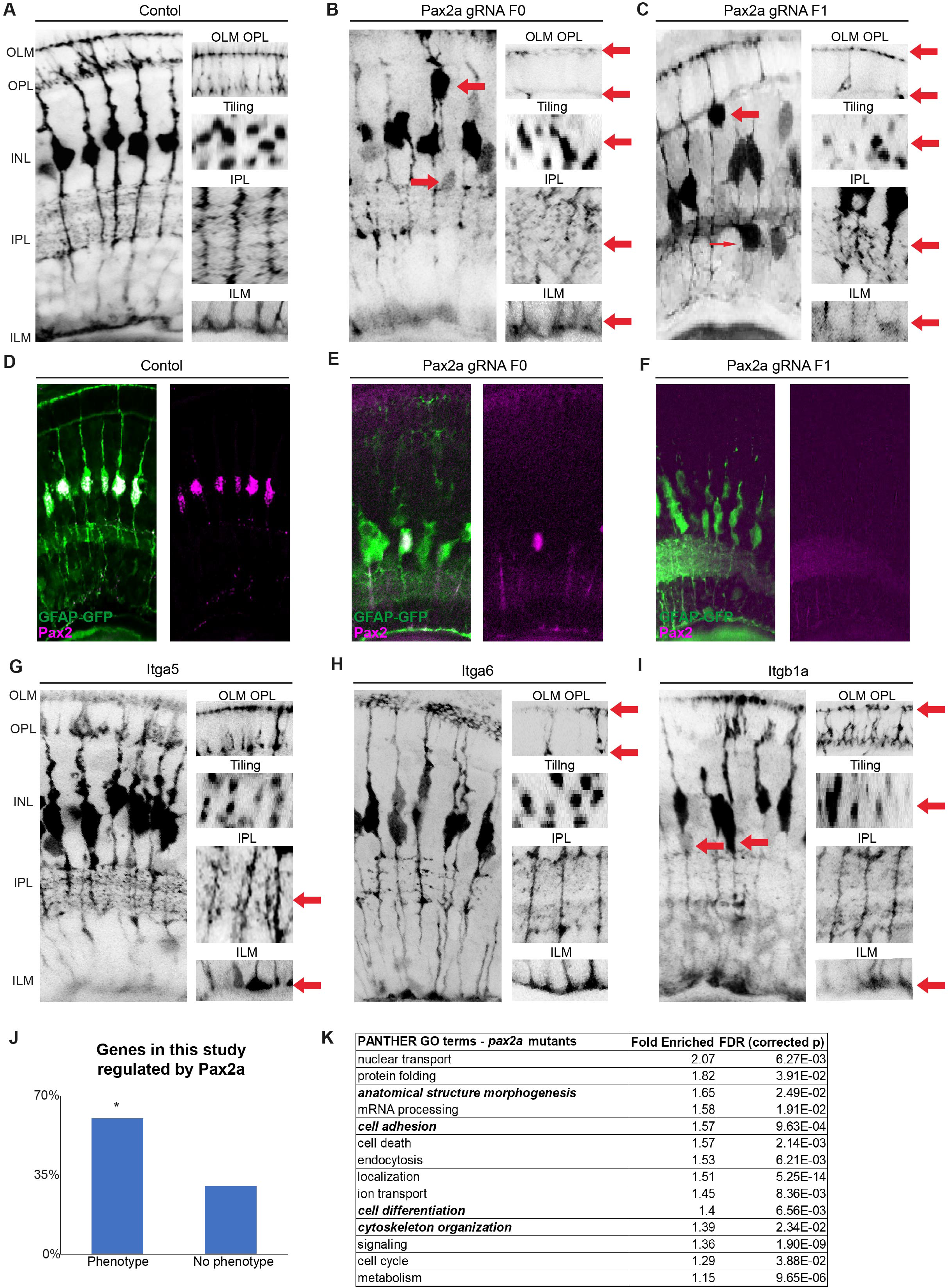
A set of highly conserved genes that affect MG cell morphology. A)*slc24a5* CRISPR injected control animals have normal MG cell morphology that extends from the apical to the basal surfaces, forming the ILM (inner limiting membrane) and OLM (outer limiting membrane) on either side.MG cells are also regularly tilled across in the eye with their cell bodies mostly restricted to the middle of the INL (inner nuclear layer) and are highly branched within the IPL (inner plexiform layer) and OPL (outer plexiform layer).B) F0 *pax2a* CRISPR injected animals have highly disorganised retinas with breaks in the OLM and ILM, abnormal tiling and apico-basal distribution of the cell bodies, as well as much less branching in the IPL and OPL.C) F1 *pax2a* CRISPR injected animals have similarly disorganised retinas, however, with more severe disruptions notable in the IPL and apico-basal distribution of cell bodies.D) In control animals (GFAP:GFP) Pax2 is expressed in all MG by 120hpf.E) F0 *pax2a* CRISPR injected animals lack Pax2 expression in most, but not all MG.F) F1 *pax2a* CRISPR injected animals Pax2 is absent from all MG.G) F0 *itga5* CRISPR injected animals have defects on the basal side of MG specifically in the ILM and IPL.H) F0 Itag6 CRISPR injected animals have defects on the apical side of the cell in the OLM and OPL.I) F0 *itb1a* CRISPR injected animals have defects in cell body tiling and apico-basal position, as well as in OLM and ILM.J) Percentages of genes used in this study that either had or did not have a phenotype.* - indicates significance by Fisher’s exact test.K) GO terms for the top 500 genes significantly (adjusted p < 0.05) up or down-regulated *pax2a* mutants.

**Figure 3.**
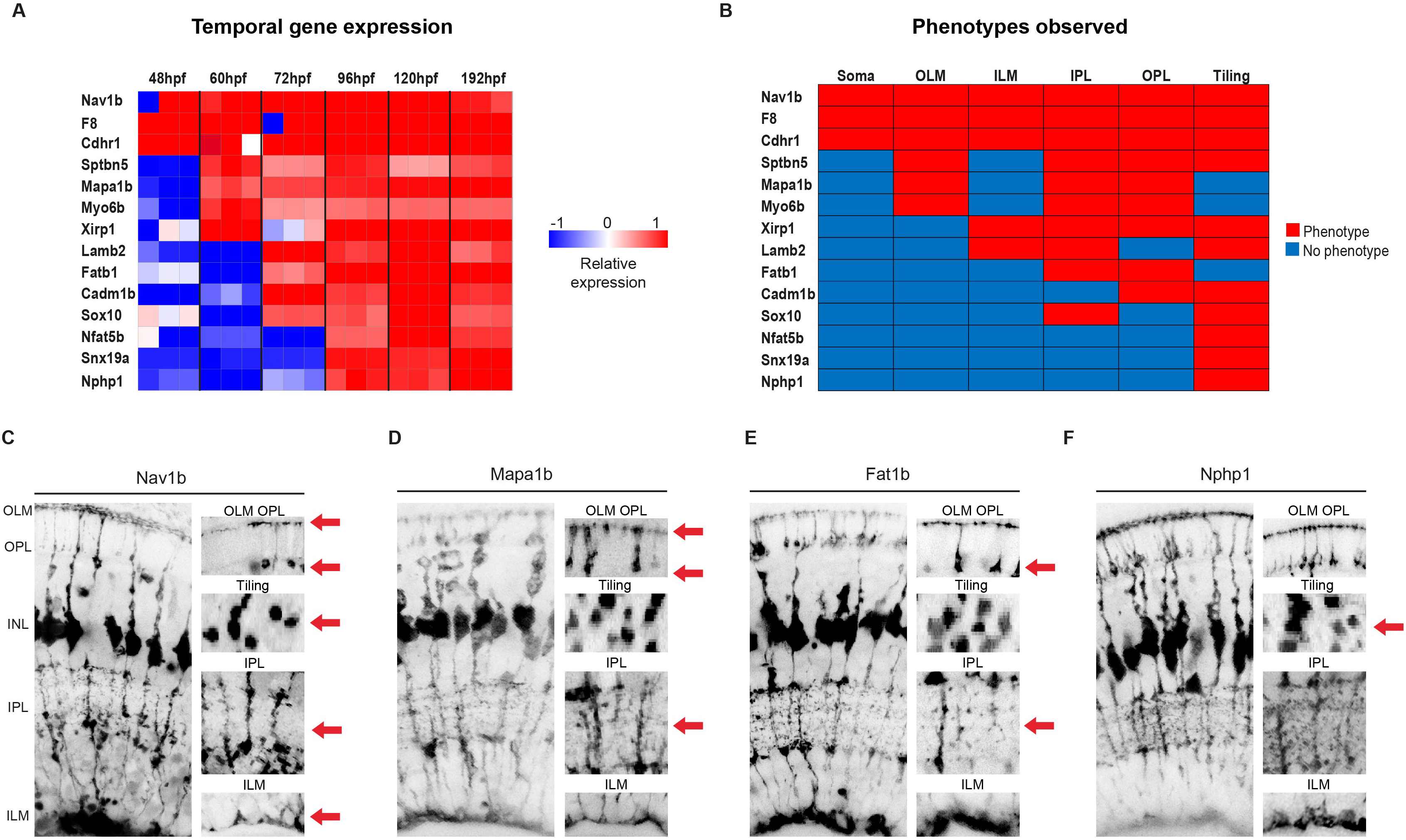
Temporal gene expression dictates MG cell morphologies. A) Heatmap to show the relative gene expression for genes tested.B) Summary of phenotypes observed for genes enriched across windows of MG cell differentiation.Red – phenotype, blue – nophenotype.C) *nav1b* CRISPR injected animals have defects in apico-basal cell body position in the INL (inner nuclear layer), OLM (outer limiting membrane), OPL (outer plexiform layer), tiling, IPL (inner plexiform layer) and ILM (inner limiting membrane).D) *mapab1* CRISPR injected animals have defects in OLM, OPL and IPL.E) *fat1b* CRISPR injected animals have defects in OPL and IPL defects.F) *nphp1* CRISPR injected animals have defects in MG cell tiling.

Overrepresentation analysis of the gene ontology GO terms identified for each developmental stage revealed dynamic changes in the biological and molecular functions of differentially expressed genes over the course of MG cell development (Extended data table 1).For example, the earliest stages (48-60hpf) of development include processes associated with cell specification, cell cycle exit, transport, growth and anatomical structure (Figure 1E).As differentiation progresses (96-192hpf), GOs of adhesion, carbohydrates and lipid metabolism, membrane transporters, and cation activity are over-represented suggesting that this is when the cells begin acquire their cell-type specific physiology (Figure 1E).

We first wanted to test whether any of the most highly conserved genes (i.e.those expressed in flies, fish and mammals) were essential for MG cell morphogenesis.Therefore, we limited our attention to those genes that code for proteins that likely impact cell morphology (e.g.cell adhesion, junction formation, cell structure or cell patterning) (Extended data table 2).We then used a CRISPR based reverse genetic screen to knock out these genes^29^ in the *Tg(GFAP:GFP)* transgenic background^30^ and looked for morphological defects in the MG cells of injected embryos at 120hpf (Extended data table 2).Control embryos injected with a guide RNA targeting the pigment gene, *slc24a5,* were devoid of much of their pigmentation while keeping MG cell shape and position unchanged (Figure 2A, Extended data fig 1C)^29^.The CRISPRed fish all continued to express the GFAP:GFP transgene in MG cells, suggesting initial glial specification is unaffected in any of the CRISPR mutants.However, we found an array of morphological defects in the MG population in many of the injected embryos.Indeed, CRISPR mutants for 11 of the 19 of these conserved genes showed defects in many of the conserved anatomical features of MG morphology (summarised in extended data table 2).Importantly, we generated F1 lines for several of these *(pax2a, nphs1, kirrela, Itga5, Itga6, wt1b, cadm1b* and *cadm4)* and confirmed that CRISPR mutation was highly specific (by DNA sequencing) and penetrant (by phenotype similarity), in agreement with previous reports (Extended data fig 2H)^29^.

One of the mutants that is particularly disruptive to MG cell morphology is in the *pax2a* gene.Pax2 is a paired domain transcription factor that has essential roles in the cellular patterning of the brain and kidney ^31^.The *pax2a* CRISPR mutation was verified by the fact that Pax2 immunostaining at 72hpf shows positive nuclei in control animals but mostly absent in F0 CRISPR injected fish and completely absent in F1 *pax2a* mutants (Figure 3D-F).CRISPR knock-outs of two nephrins *(nphs1* and *kirrela)* and the transcription factor Wt1 b, all of which are known to work with Pax2 during kidney and CNS differentiation ^31–33^ also resulted in defects in many MG cell morphological features (Extended data fig 2A-C).In the kidney podocytes, the expression of membrane spanning Nephrins are linked to the expression of Pax2 and Wt1 ^31^, and it is interesting to note that the two *nephrin* mutants have more limited defects and than either *pax2a* or *wt1b* mutants.Remarkably, analysis of the transcriptome of MG cells in *pax2a* mutants shows that 60% of the genes we that we tested that affect MG cell morphogenesis have significant changes of expression, whereas only about 30% of those that did not affect MG cell morphogenesis were misregulated in *pax2a* mutants (Figure 2J).Furthermore, GO analysis on the differentially expressed genes in Pax2a mutant MG cells shows a significant enrichment of GO terms related to cell morphology, adhesion and differentiation (Figure 1K).These results suggest that Pax2a may be a “master regulator” of MG cell morphology (Supp table 3).

The rapid co-ordinated expansion of MG cell morphological domains along the apical and basal limiting membranes and within the cellular layers of the retina suggests differential interactions with the extracellular matrix (ECM) and neurons are likely to be involved in MG cell morphogenesis.Indeed, the ECM receptor, integrin has previously been associated with glial radial glial defects ^34^, and we found that several of these *integrin* genes when mutated give rise to morphological phenotypes in MG cells (Figure 2G-I).It is particularly interesting that different members of the integrin family affect different morphological features of MG cells.Mutants for *itga5* have defects in the ILM and IPL (Figure 2G), *itga6* mutants have defects in the OLM, OPL (Figure 2H), and *itgB1a* mutants have defects in apical-basal positioning as well as in the OLM and ILM (Figure 2I).Mmp2 is a critical modulator of the ECM ^35–38^, and we found that *mmp2* mutants share defects with these integrin mutants (Extended data figure 2D).Mutants of the cell adhesion molecules Cadm4, Cadm1a and Cadm2b, in addition to unshared specific defects the plexiform layers, all show irregular MG cell tiling (Extended data figure 2E - G).Although the focus here is on those genes that are expressed in MG cells themselves, the involvement of such adhesion molecules, also fuels the understanding that MG cell morphogenesis cannot be an entirely autonomous process.MG cells must shape themselves appropriately with respect to their neighbours within the tissue of the retina.It would this be interesting to look for non-autonomous factors such as the ligands for these adhesion molecules.

The fact that such a large fraction of the conserved genes we tested affected MG cell morphogenesis, allowed us to ask whether there is any functional correlation between the temporal expression of morphogenetic genes and the sequentially arising features of these cells.We first used hierarchical clustering and Trimmed Means of M (TMM) differential expression analysis, to identify lists of genes that first became enriched at specific time points and remained enriched until 192hpf (i.e.48-192hpf, 60-192hpf, 72-192hpf and 96-192hpf) (supplemental table 3).Then, we again used CRISPR/Cas9 to knock down candidates associated with cellular differentiation, cell adhesion, cell morphology and cell dynamics (Extended data table 2).Genes that were enriched from 48-192hpf included *nav1b* (part of the neuron navigator family of genes with limited recognised functions), *f8* (a coagulation factor exclusively known for its involvement for blood clotting), and *cdhr1* (a protocadherin which is highly expressed in the mature retina and link to human retinitis pigmentosa and cone-rod dystrophy) ^39^ (Figure 3A).All of these, when mutated, produced highly irregularly shaped MG cells with defects in many of the cells conserved morphological features (Figure 3B, C,; Extended data figure 3B, C,).Genes enriched from 60-192 hpf with mutant phenotypes include: *sptbn5* (a beta-spectrin that forms part of the cytoskeleton), *myo6b* (a myosin motor protein), *xirp1* (an actin-binding protein associated with adherens junctions), and *map1ap* (a microtubule-associated protein) (Figure 3A).Each of these has defects in multiple, yet specific, aspects of MG cell morphogenesis but none showed apico-basal positioning defects (Figure 3B, D; Extended data figure 3D-H).Genes enriched from 72-192hpf with mutant phenotypes include: *lamb2* (laminin component of extracellular matrix), *fat1b* (an a-typical cadherin involved in Hippo signalling, *cadm1b* (also part of the cell adhesion molecule family), and *sox10* (SOX family transcription factor associated with peripheral glial differentiation) (Figure 3A).These displayed defects in fewer aspects of MG cell morphogenesis without defects in apico-basal positioning or OLM formation (Figure 3B, E; Extended data figure 3G – I).Genes enriched from 96-192hpf with mutant phenotypes include: *nfat5c* (a transcription factor associated with osmotic stress), *snx19a* (sorting nexin associated with g-protein coupled signalling), *nphp1* (a nephrocystin thought to be associated cilia-mediated signalling) (Figure 1A).All of these only showed defects in MG cell tiling (Figure 3B, F; Extended data figure 4 J, K).This gradation of phenotypes with genes enriched at early stages causing multiple severe defects in MG cell morphogenesis and later enriched genes having milder and fewer phenotypes suggest the reasonable hypothesis that the basic that some genes are involved in many steps of cellular morphogenesis, while others, especially those expressed later are involved in fewer steps.

**Figure 4.**
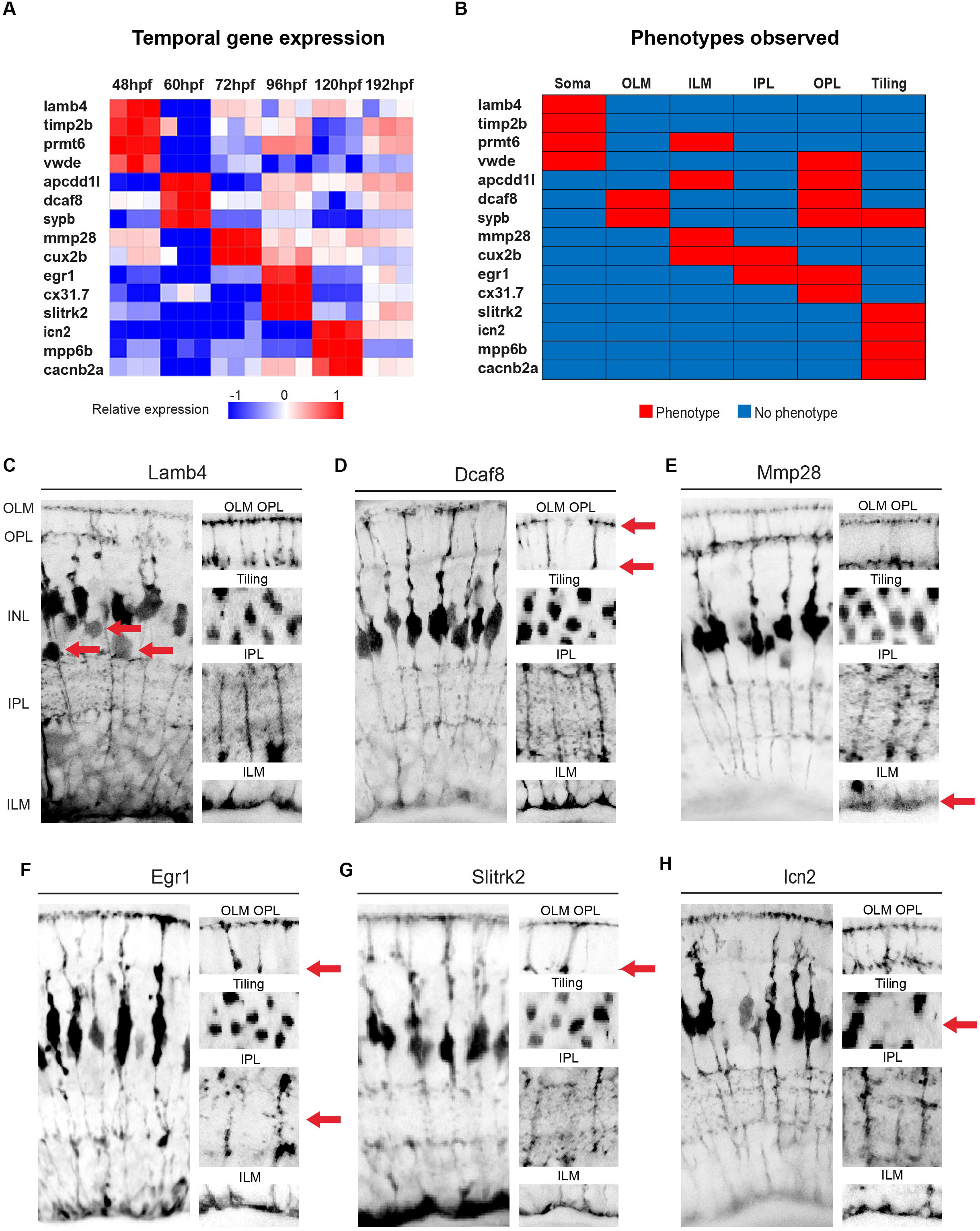
Discrete gene expression regulates MG cell compartment morphology. A) Heatmap to show the relative gene expression for genes tested.These were all screen in F0 CRISPR injected mutants.B) Summary of phenotypes observed for genes enriched across windows of MG differentiation.Red – phenotype, blue – no-phenotype.C) *lamb4* mutants have defects in apico-basal distribution of MG cell bodies only.D) *dcaf8* mutants have defects in the OLM and OPL.E) *mmp28* CRISPR injected mutants have defects in the ILM only.F) *egr1* mutants have defects in the IPL and OPL.G) *slitkr2* mutants have defects in the OPL layer only.

To explore the temporal correlation further, we tested candidate genes whose transcripts were enriched during specific time points of development (Figure 4A).When we target two ECM proteins enriched at 48hpf, Lamb4 (laminin) and Timp2b (an inhibitor of Mmps), we observed defects in apical-basal soma positions (Figure 4B, C; Extended data figure 4B), consistent with the fact that soma positioning occurs first between 48hpf and 60hpf (Figure 1B).Apical-basal soma defects were also seen in mutants of two other genes that were transiently enriched at 48hpf: *prtm6a* (a nuclear methylase) and *vwde* (Von Willebrand blood coagulation factor) (Figure 4B; Extended data figure 4C, D).In addition, *prmt6* mutants show defects in ILM formation, and *vwde* mutants show defects in OPL formation (Figure 4B, Extended data figure 4C, D).When we targeted genes that were overrepresented at 60hpf, we observed no defects in apical-basal somal positioning, yet we did find defects in later stages of morphogenesis (Figure 4B).For instance, *dcaf8* (unknown molecular function) mutants have OPL and OLM defects (Figure 4D), *apcdd1l* (unknown molecular function) mutants have defects in OPL and ILM defects and MG tiling (Extended data figure 4E), and *sypb* (a synaptic vesicle-associated protein) have MG spacing, OPL, OLM defects (Extended data figure 4F).Genes enriched at 72hpf include *mmp28* (a protease regulator of extracellular matrix) and *cux2b* (a Cut-like transcription factor) (Figure 4A).These mutants had a more limited set of effects only in the ILM of *mmp28* mutants (Figure 4E) and effects on ILM and IPL in *cux2b* mutants (Figure 4B; Extended data figure 4G).Genes enriched at 96hpf include *cx31.7* (a gap junction connexin), *egr1* (a zinc finger transcription factor), and *slitrk2* (an integral membrane protein) (Figure 4A).*egr1* mutants have subtle defects in the IPL and OPL (Figure 4F), *slitrk2* mutants only have defects in the OPL (Figure 4G), and *cx31.7* only show tiling defects (Extended data figure 4H).At 120hpf *icn2* (a gene of unknown molecular function), *mpp6b* (a membrane-associated guanylate kinase) and *cacnb2a* (a subunit voltage-dependent calcium channels) are enriched (Figure 4A).These mutants produce nothing more than MG cell tiling defects, (Figure 4H; Extended data figure 4G).Together, these data show a correspondence between the temporal patterning of gene expression and the development of particular features of cellular anatomy, suggesting an approach to begin to dissect the developmental history of the cellular morphogenesis.

The high level of conserved gene expression in glial cells within the animal kingdom, especially of those genes involved in MG cell morphogenesis, suggests that cell-specific morphogenetic processes are likely to be broadly shared.Many of these genes (or their homologues) are known from previous work to be involved in various aspects of cellular morphogenesis.The clearest example of this is the relationship between the *Pax2* and *Wt1* genes that have been identified as crucial for cellular patterning through their regulation of the Nephrins in kidney development in vertebrates ^31–33^ and for cell shape and patterning of glia in the fly eye ^40,41 28^^42^–^44^.The fact that the expression 60% of our candidate genes with essential roles in MG morphogenesis are regulated by Pax2a suggests a hierarchy of genetic regulation leading to effector genes such as the nephrins and integrins that carry out specific morphogenetic roles.It is thus noteworthy that three integrins (Itga1a, Itga1b and Itgb1a), which have been shown to play a role in radial glial morphogenesis in the cortex ^45,46^, are each required for distinct features of MG cell morphology (Figure 2).This idea is strengthened by our finding of the expression of two beta-laminins is enriched in MG cells.*lamb4* is enriched early on (48hpf) and effects apical-basal soma position while *lamb2* is enriched from 60hpf onwards and is required in the spacing of MG cells as well as IPL formation (Figure 4).We also found some unexpected genes with roles in MG cell morphogenesis.Two of these include *vwf* and *f8* which have been extensively studied in the context of blood coagulation, and both have known human mutants that lead to human bleeding diseases ^47^.In MG cells, both factors are expressed early in differentiation and have a severe effect on MG shape and spacing.

Our results point to specific genetic repertoires working at particular periods of development to generate specific anatomical features of MG cell morphogenesis.The fact that many of the genes we identified are conserved during MG cell development across the animal kingdom indicates that temporally conserved genetic programmes of cell shape and patterning evolved early.We believe that many of the principles of developmental biology gathered from the study of multicellular organic morphogenesis may be relevant to in the further studies of cellular morphogenesis, and that this study may provide an entry point for the further dissection of the molecular mechanisms of cell morphogenesis.

## Methods

### Animals

Adult zebrafish were maintained and bred at 26.5°C.Embryos were raised at 25°C– 32°C and staged based on hpf ^48^ Embryos were treated with 0.003% phenylthiourea (Sigma) from 10 hpf to prevent pigmentation.All animal work was approved by Local Ethical Review Committee at the University of Cambridge and performed according to the protocols of project license PPL 80/2198.

### Transgenic Lines

Transgenic lines Tg*(atoh7:gap43-mRFP1)cu2^49^, Tg(GFAP:GFP)^30^, Tg(TP1:Venus-Pest) ^12^*.

### FACS, RNA-seq and Bioinformatics

20-40 whole eyes of *Tg(GFAP:GFP)* fish were dissected from each developmental time point (48, 60, 72, 96, 120 and 192 hpf) in and washed several times to remove debris in L-15 (Leibovitz’s L-15 Medium).Eyes were then incubated in Trypsin-EDTA 0.25% (Sigma) at 37*C for 15min, washed several times and dissociated using FBS coated pipette tips in Calcium-free medium (116.6 mM NaCl, 0.67 mM KCl, 4.62 mM Tris, 0.4 mM EDTA).Single cell suspensions were sorted on a Beckman Coulter MoFlo to capture Muller glia (GFP) and control retinal tissue (non-GFP).Cells were sorted into lysis buffer, and RNA was immediately extracted using the RNeasy mini kit (Qiagen).RNA concentration and qualities were assessed on an Agilent Bioanalyzer and RNA amplification and cDNA synthesis was performed with the Ovation RNA Amplification System V2 (NuGEN) using manufacturer’s protocol.Nextera library preparations were performed using the Nextera DNA library kit according to the manufacturer’s directions and sent to the Sanger Center for sequencing.

Sequence files were paired, trimmed and aligned using Hisat2 to the zebrafish genome (version: Zv9) and RNA-seq bioinformatic and statistical analysis was performed in R using the Bioconducter, Featurecounts, Rsubread, limma, DESeq2, DEFormats, pheatmap, ggplots, org.Dr.er.db, and EdgeR packages.Cross-species gene conversions were performed using Ensembl (Biomart) and statistical significance of gene overlaps was done using a Fisher’s exact test with Bonferroni correction.Gene Ontology analysis and statistics were performed using Gene Ontology Consortium ^50,51^.

### Embryo Manipulations

For blastomere transplantations, high-to oblong-stage embryos were dechorionated by pronase digestion (Sigma), placed in agarose moulds, and between 5 and 30 blastomeres were transferred between *Tg(TP1:Venus)* embryos to wildtype embryos using a glass capillary connected to a 2 ml syringe.Embryos were grown on dishes coated with 1% agarose in 0.04% PTU overnight until imaged by confocal microscopy.

### sgRNA design and Reverse Genetic Screen

The sgRNA design and strategy are largely based on the methods from Shah and colleagues ^29^.Briefly, each guide RNA was designed using the ChopChop design tool ^52^ at chopchop.cbu.uib.no/index.php.For each gene, the two gRNAs with minimal predicted off-target sites were selected.In the first screen these we picked the targets with overall ranking while in the second screen we used the highest ranking targets for the first and last exons of each gene.Template DNA was synthesised by *in vitro* transcription of a two oligo PCR method.For this, an oligo scaffold containing the RNA loop structure

5’[gatccgcaccgactcggtgccactttttcaagttgataacggactagccttattttaacttgctatttctagctctaaaac ]3’ required for Cas9 was synthesised and used for the syntheses of all gRNAs (Extended data table 2).Next, a unique oligo containing the T7 promoter, the 20 nucleotides gRNA, and 20 bases of homology to the scaffold oligo was synthesised.PCR amplification of these annealed oligos sequence was created using Phusion master mix (England BioLabs, M0531L) with 10uM scaffold and gRNA for 40 cycles in a thermal cycler.This PCR product was purified (PCR purification kit - Qiagen) and used as a template for the *in vitro* transcription reaction (T7 megascript – Ambion).RNA was purified on columns (Zymo Research, D4014) and injected using 100pg of each gRNA (200ng total) with 1200pg of Cas9 encoding mRNA.

### Immunostaining, Microscopy and data analysis

For immunostaining samples were fixed in 4% paraformaldehyde overnight at 4°C, washed in PBS and then stored in MeOH at -20°C until used.Samples were rehydrated in a MeOH:PBS series (3:1, 1:1, 1:3) followed by three PBST (PBS + 0.05% Triton-X100) washes.Primary and secondary antibodies were diluted in PBS using the following concentrations: Rabbit anti-Pax2 1:200 (Sigma), goat anti-rabbit conjugated Alexa Fluor 555 1:500 (Invitrogen) and GFP-Booster Atto488 1:500 (Chormotek).Samples were mounted on slides with a coverslip bridge (to prevent crushing the tissue) in Prolong Diamond (Invitrogen) and allowed to cure at room temperature overnight before imaging.

Laser scanning confocal imaging was performed using an Olympus FV1000 microscope with a 60 X oil objective (1.35 NA).For live imaging, optical sections at 0.5–1 μm separation were taken to cover the region of the retina containing the cells of interest (between 40 and 100μm) every 15 minutes over a 12 hour period.Confocal imaging of live and fixed embryos was performed as described previously ^53^ Confocal data was analysed and processed using Volocity (Improvision) and ImageJ/FIJI (NIH).

**Extended Data table 1: Gene ontologies of each stage of MG cell differentiation.**

**Extended Data table 2: Summary of all genes used with gRNAs and phenotypes.**

**Supplemental table 1: TMM fold change analysis of each MG cell developmental stage.**

**Supplemental table 2: Log counts per million for all samples**

**Supplemental table 3: TMM gene enrichments of Pax2a mutant MG cells**

---

**Extended Data Figure 1.**
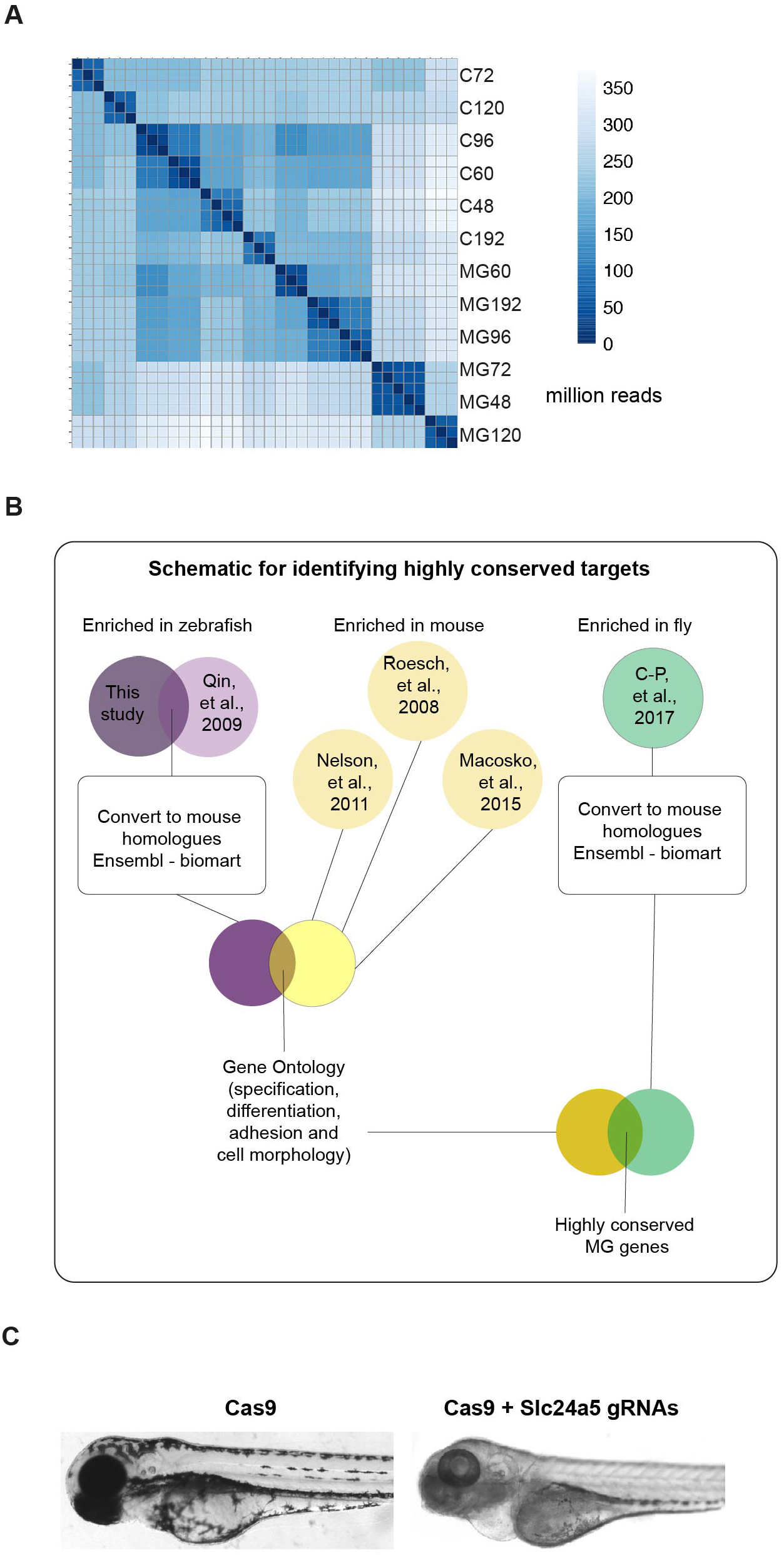
Temporal genetics of MG cell differentiation. A) Hierarchical clustering of samples used for RNA-seq demonstrating consistency between the three replicates used for each time point (MG-GFAP-GFP sorted cells, C - GFP negative control tissue).B) Schematic representation of how highly conserved genes we bioinformatically identified.C) Cas9 only injected fish have normal pigmentation at 120hpf while those injected with Cas9 and the slc45a5 guide RNAs are mostly devoid of pigment.

**Extended Data Figure 2.**
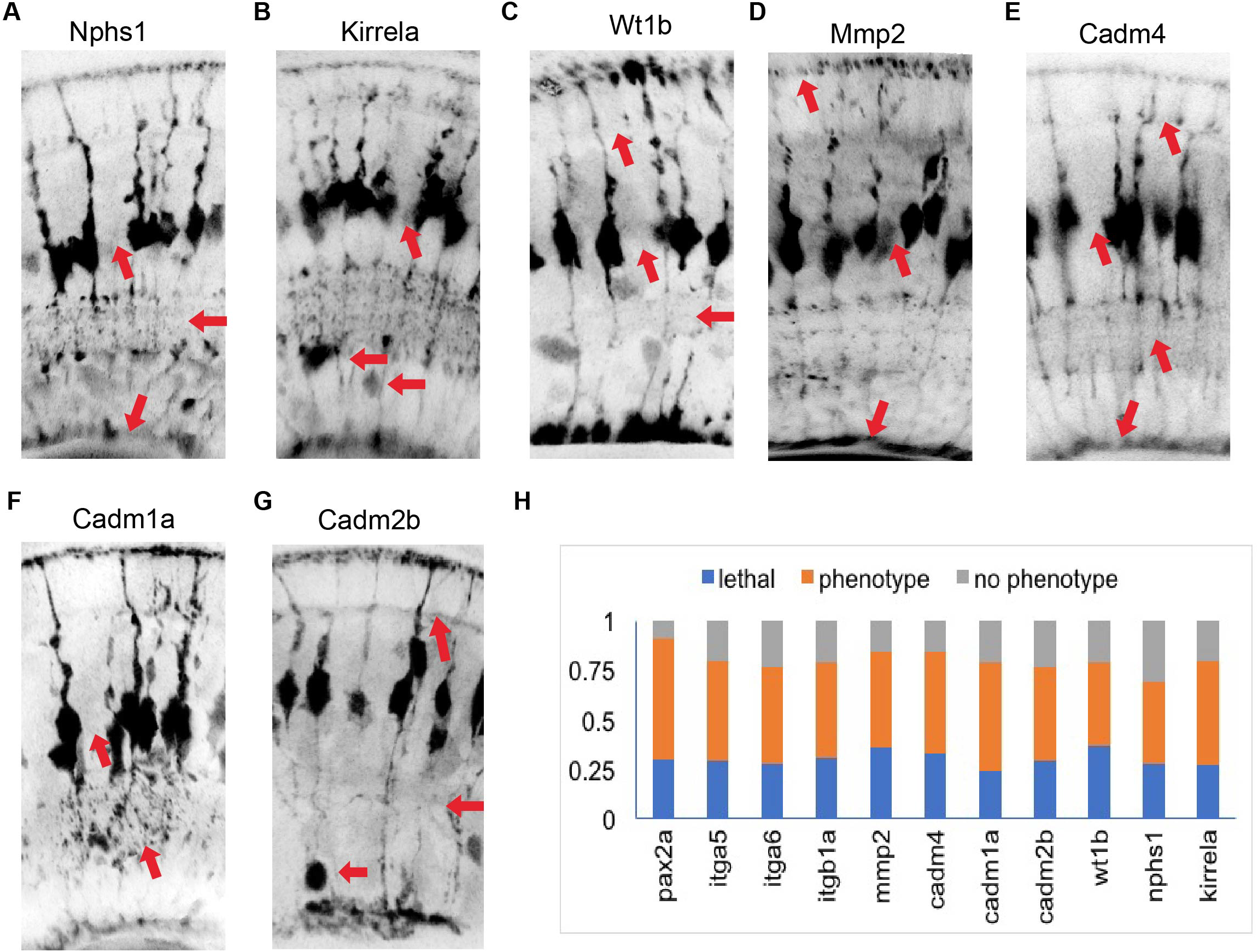
Phenotypes of conserved highly conserved MG cell genes. A) mmp2 mutants have defects in OLM, ILM and tiling.B) cadm4 mutants have defects in OPL, IPL, ILM and tiling.C) Cadm1a mutants have defects in IPL and tiling.D) cadm2b mutants have defects in a cell body positing, IPL OPL and tiling.E) wt1 mutants have defects in cell body position, IPL, OPL and tiling.F) nphs1 mutants have defects in ILM, IPL and tiling.G) kirrela mutants have defects in cell body position and tiling.H) Proportions of injected animals that died, had no phenotype or had a phenotype after injection by 120hpf.

**Extended Data Figure 3.**
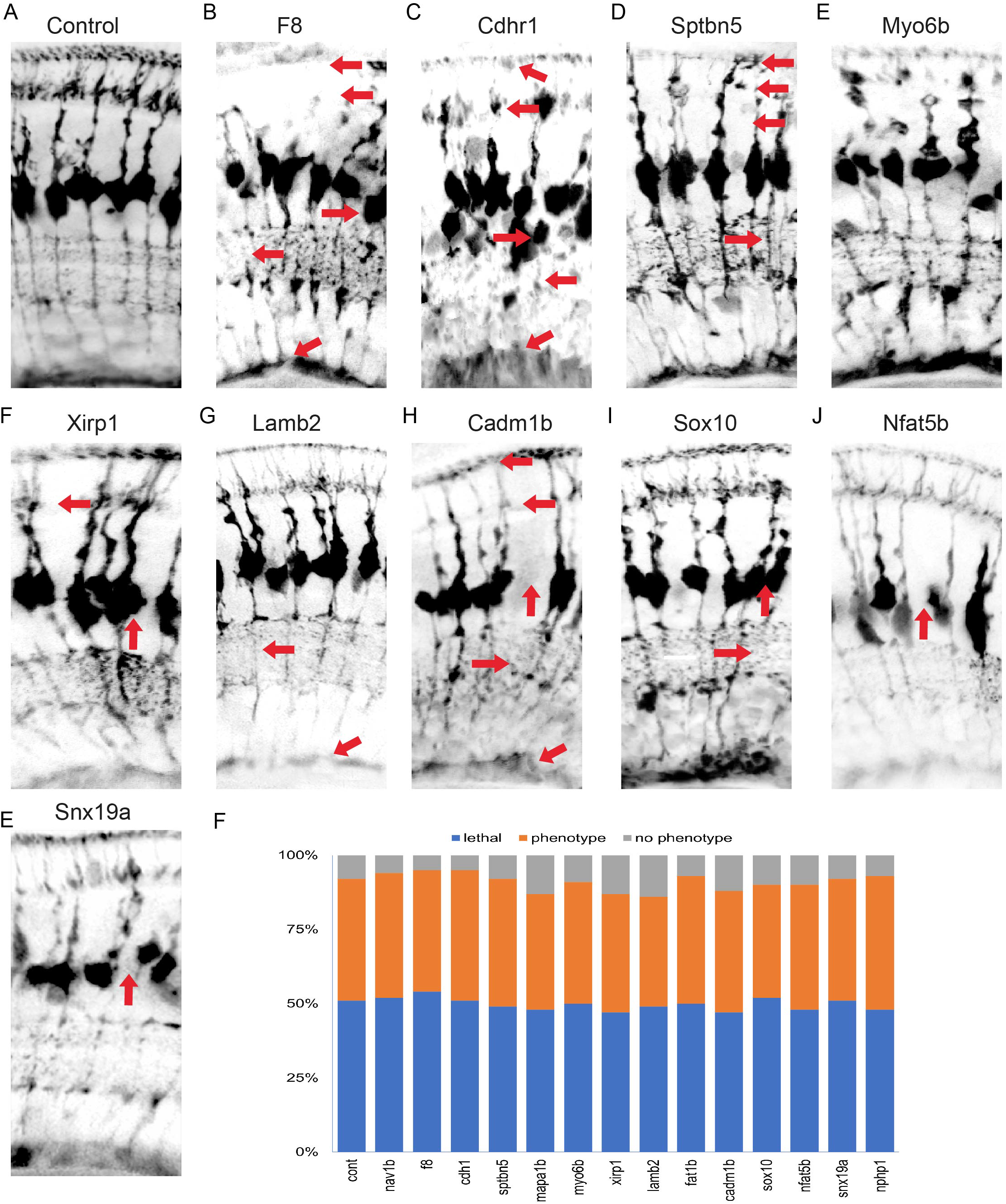
Phenotypes of gene mutants enriched over windows of MG cell differentiation. A) slc45a5 controls have no observable MG phenotype.B) f8 mutants have defects in cell body position, OLM, ILM, IPL, OPL and tiling.C) cdhr1 mutants have defects in cell body position, OLM, ILM, IPL, OPL and tiling.D) sptbn mutants have defects in OLM, IPL, OPL and tiling.E) mapa1b mutants have defects in OLM, IPL and OPL.F) xirp1 mutants have defects in OPL and tiling.G) lamb2 mutants have defects in ILM, IPL and tiling.H) Cadm1 b mutants have defects in ILM, IPL, OPL and tiling.I) sox10 mutants have defects in IPL and tiling.J) nfat5 mutants have tiling defects.K) snx19a mutants have tiling defects.L) Proportions of injected animals that died, had no phenotype or had a phenotype after injection by 120hpf.

**Extended Data Figure 4.**
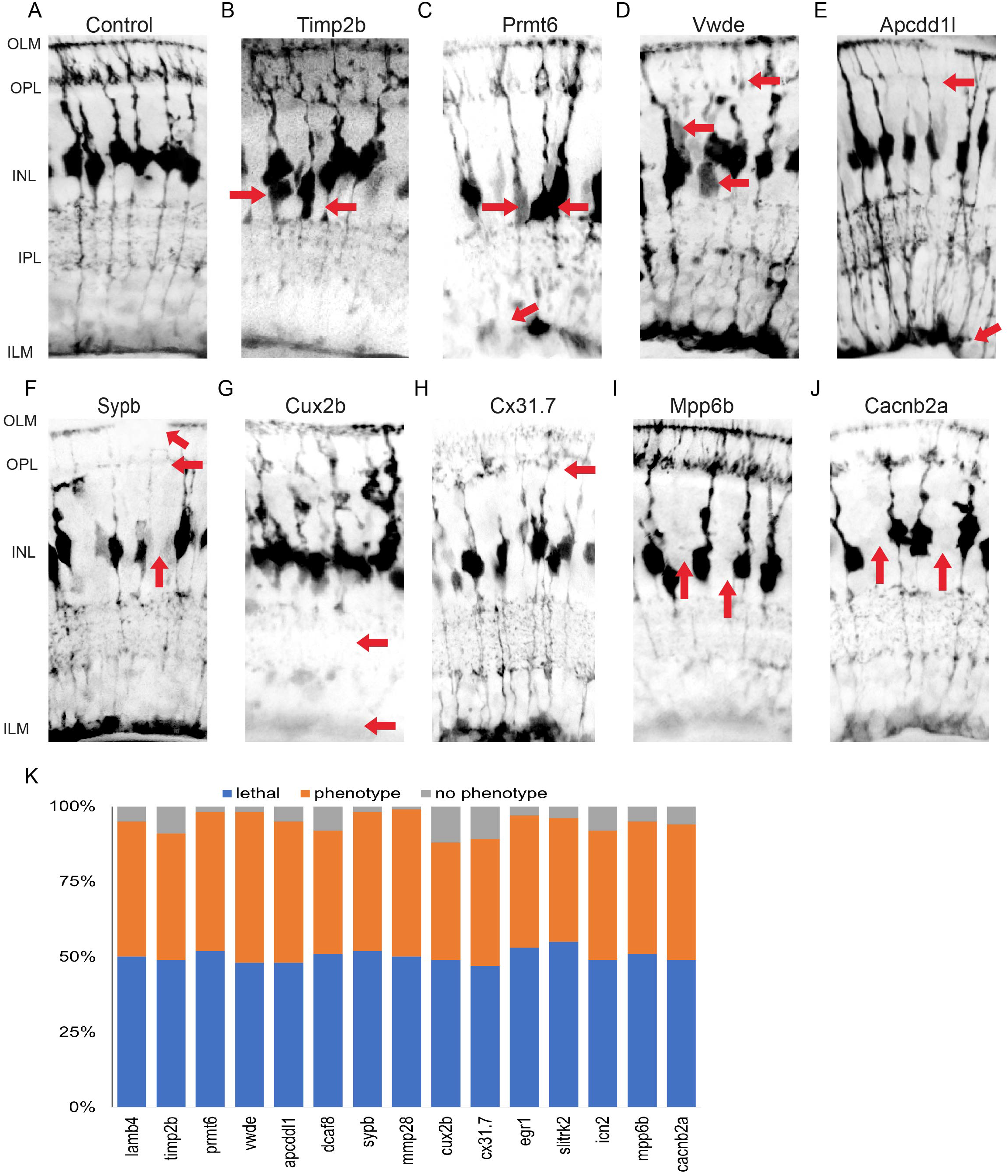
Phenotypes of gene mutants that are enriched at specific times of MG differentiation. A) slc45a5 controls have no observable MG phenotype.B) timp2b mutants have defects in cell body position.C) prmt6 mutants have defects in cell body position and ILM.D) vwde mutants have defects in cell body position, OLM, ILM, IPL, OPLand tiling.E) apcdd1l mutants have defects in IPL and OPL.F) sypb mutants have defects in OLM, OPL and tiling.G) Cux2b mutants have defects in ILM and IPL.H) cx31.7 mutants have defects in tiling.I) Mpp6b mutants have defects in tiling.J) cacnb2a mutants have defects in tiling.H) Proportions of injected animals that died, had no phenotype or had a phenotype after injection by 120hpf.

## References

1. Kandel, E. Principles of Neural Science, Fifth Edition. (McGraw Hill Professional, 2013).

2. Muller & H. Zur Histologie der Netzhaut. Zeitschift fur Wissenschaft und Zoologie 3, 234–237 (1851).

3. Cajal, S.R.y. La rétine des vertébrés. (1892).

4. Jadhav, A.P., Roesch, K. & Cepko, C.L. Development and neurogenic potential of Müller glial cells in the vertebrate retina. Prog.Retin.Eye Res. 28, 249–262 (2009).

5. Reichenbach, A. & Bringmann, A. New functions of Müller cells. Glia 61, 651–678 (2013).

6. MacDonald, R.B., Charlton-Perkins, M. & Harris, W.A. Mechanisms of Müller glial cell morphogenesis. Curr.Opin.Neurobiol. 47, 31–37 (2017).

7. MacDonald, R.B. et al. Müller glia provide essential tensile strength to the developing retina. J.Cell Biol. 210, 1075–1083 (2015).

8. Williams, P.R. et al. In vivo development of outer retinal synapses in the absence of glial contact. J.Neurosci. 30, 11951–11961 (2010).

9. Wang, J. et al. Anatomy and spatial organization of Müller glia in mouse retina. J.Comp.Neurol. 525, 1759–1777 (2017).

10. Kolb, H., Nelson, R., Ahnelt, P. & Cuenca, N. Cellular organization of the vertebrate retina. Prog.Brain Res. 131, 3–26 (2001).

11. Reichenbach, A. & Reichelt, W. Postnatal development of radial glial (Müller) cells of the rabbit retina. Neurosci. Lett. 71, 125–130 (1986).

12. Ninov, N., Borius, M. & Stainier, D.Y.R. Different levels of Notch signaling regulate quiescence, renewal and differentiation in pancreatic endocrine progenitors. Development 139, 1557–1567 (2012).

13. Bushong, E.A., Martone, M.E., Jones, Y.Z. & Ellisman, M.H. Protoplasmic astrocytes in CA1 stratum radiatum occupy separate anatomical domains. J.Neurosci. 22, 183–192 (2002).

14. Biehlmaier, O., Neuhauss, S.C.F. & Kohler, K. Synaptic plasticity and functionality at the cone terminal of the developing zebrafish retina. J.Neurobiol. 56, 222–236 (2003).

15. Eisenfeld, A.J., Bunt-Milam, A.H. & Sarthy, P.V. Müller cell expression of glial fibrillary acidic protein after genetic and experimental photoreceptor degeneration in the rat retina. Invest.Ophthalmol.Vis.Sci. 25, 1321–1328 (1984).

16. Lehre, K.P., Davanger, S. & Danbolt, N.C. Localization of the glutamate transporter protein GLAST in rat retina. Brain Res. 744, 129–137 (1997).

17. White, R.D. & Neal, M.J. The uptake of L-glutamate by the retina. Brain Res. 111, 79–93 (1976).

18. Saari, J.C. et al. Visual cycle impairment in cellular retinaldehyde binding protein (CRALBP) knockout mice results in delayed dark adaptation. Neuron 29, 739–748 (2001).

19. Jo, A.O. et al. TRPV4 and AQP4 Channels Synergistically Regulate Cell Volume and Calcium Homeostasis in Retinal Müller Glia. J.Neurosci. 35, 13525–13537 (2015).

20. Dahlin, A., Royall, J., Hohmann, J.G. & Wang, J. Expression profiling of the solute carrier gene family in the mouse brain. J.Pharmacol.Exp.Ther. 329, 558–570 (2009).

21. Zong, H. et al. Hyperglycaemia-induced pro-inflammatory responses by retinal Müller glia are regulated by the receptor for advanced glycation end-products (RAGE). Diabetologia 53, 2656–2666 (2010).

22. Riepe, R.E. & Norenburg, M.D. Müller cell localisation of glutamine synthetase in rat retina. Nature 268, 654–655 (1977).

23. Lehre, K.P. & Danbolt, N.C. The number of glutamate transporter subtype molecules at glutamatergic synapses: chemical and stereological quantification in young adult rat brain. J.Neurosci. 18, 8751–8757 (1998).

24. Qin, Z., Barthel, L.K. & Raymond, P.A. Genetic evidence for shared mechanisms of epimorphic regeneration in zebrafish. Proc.Natl.Acad.Sci.U.S.A. 106, 9310–9315 (2009).

25. Roesch, K. et al. The transcriptome of retinal Müller glial cells. J.Comp.Neurol. 509, 225–238 (2008).

26. Nelson, B.R. et al. Genome-wide analysis of Müller glial differentiation reveals a requirement for Notch signaling in postmitotic cells to maintain the glial fate. PLoS One 6, e22817 (2011).

27. Macosko, E.Z. et al. Highly Parallel Genome-wide Expression Profiling of Individual Cells Using Nanoliter Droplets. Cell 161, 1202–1214 (2015).

28. Charlton-Perkins, M.A., Sendler, E.D., Buschbeck, E.K. & Cook, T.A. Multifunctional glial support by Semper cells in the Drosophila retina. PLoS Genet. 13, e1006782 (2017).

29. Shah, A.N., Davey, C.F., Whitebirch, A.C., Miller, A.C. & Moens, C.B. Rapid Reverse Genetic Screening Using CRISPR in Zebrafish. Zebrafish 13, 152–153 (2016).

30. Bernardos, R.L. & Raymond, P.A. GFAP transgenic zebrafish. Gene Expr.Patterns 6, 1007–1013 (2006).

31. Quaggin, S.E. Transcriptional regulation of podocyte specification and differentiation. Microsc.Res.Tech. 57, 208–211 (2002).

32. Ambu, R. et al. WT1 expression in the human fetus during development. Eur.J.Histochem. 59, 2499 (2015).

33. Putaala, H., Soininen, R., Kilpeläinen, P., Wartiovaara, J. & Tryggvason, K. The murine nephrin gene is specifically expressed in kidney, brain and pancreas: inactivation of the gene leads to massive proteinuria and neonatal death. Hum.Mol.Genet. 10, 1–8 (2001).

34. Georges-Labouesse, E., Mark, M., Messaddeq, N. & Gansmüller, A. Essential role of alpha 6 integrins in cortical and retinal lamination. Curr.Biol. 8, 983–986 (1998).

35. Yamana, S. et al. The Cell Adhesion Molecule Necl-4/CADM4 Serves as a Novel Regulator for Contact Inhibition of Cell Movement and Proliferation. PLoS One 10, e0124259 (2015).

36. Moiseeva, E.P., Straatman, K.R., Leyland, M.L. & Bradding, P. CADM1 controls actin cytoskeleton assembly and regulates extracellular matrix adhesion in human mast cells. PLoS One 9, e85980 (2014).

37. Turksen, K., Opas, M. & Kalnins, V.I. Cytoskeleton, Adhesion, and Extracellular Matrix of Fetal Human Retinal Pigmented Epithelial Cells in Culture. Ophthalmic Res. 21, 56–66 (1989).

38. Li, D. et al. CADM2, as a new target of miR-10b, promotes tumor metastasis through FAK/AKT pathway in hepatocellular carcinoma. J.Exp.Clin.Cancer Res. 37, 46 (2018).

39. Stingl, K. et al. CDHR1 mutations in retinal dystrophies. Sci.Rep. 7, 6992 (2017).

40. Cagan, R. The signals that drive kidney development: a view from the fly eye. Curr.Opin.Nephrol.Hypertens. 12, 11–17 (2003).

41. Bao, S. & Cagan, R. Preferential Adhesion Mediated by Hibris and Roughest Regulates Morphogenesis and Patterning in the Drosophila Eye. Dev.Cell 8, 925–935 (2005).

42. Flores, G.V., Daga, A., Kalhor, H.R. & Banerjee, U. Lozenge is expressed in pluripotent precursor cells and patterns multiple cell types in the Drosophila eye through the control of cell-specific transcription factors. Development 125, 3681–3687 (1998).

43. Fu, W. & Noll, M. The Pax2 homolog sparkling is required for development of cone and pigment cells in the Drosophila eye. Genes Dev. 11, 2066–2078 (1997).

44. Charlton-Perkins, M. et al. Prospero and Pax2 combinatorially control neural cell fate decisions by modulating Ras-and Notch-dependent signaling. Neural Dev. 6, 20 (2011).

45. Belvindrah, R., Graus-Porta, D. & Goebbels, S. β1 integrins in radial glia but not in migrating neurons are essential for the formation of cell layers in the cerebral cortex. Journal of (2007).

46. Förster, E. et al. Reelin, Disabled 1, and beta 1 integrins are required for the formation of the radial glial scaffold in the hippocampus. Proc.Natl.Acad.Sci.U.S.A. 99, 13178–13183 (2002).

47. Desch, K.C. Regulation of plasma von Willebrand factor. F1000Res. 7, 96 (2018).

48. Kimmel, C.B., Ballard, W.W., Kimmel, S.R., Ullmann, B. & Schilling, T.F. Stages of embryonic development of the zebrafish. Dev.Dyn. 203, 253–310 (1995).

49. Zolessi, F.R., Poggi, L., Wilkinson, C.J., Chien, C.-B. & Harris, W.A. Polarization and orientation of retinal ganglion cells in vivo. Neural Dev. 1, 2 (2006).

50. Ashburner, M. et al. Gene ontology: tool for the unification of biology. The Gene Ontology Consortium. Nat.Genet. 25, 25–29 (2000).

51. The Gene Ontology Consortium. Expansion of the Gene Ontology knowledgebase and resources. Nucleic Acids Res. 45, D331–D338 (2017).

52. Labun, K., Montague, T.G., Gagnon, J.A., Thyme, S.B. & Valen, E. CHOPCHOP v2: a web tool for the next generation of CRISPR genome engineering. Nucleic Acids Res. 44, W272–6 (2016).

53. Das, T., Payer, B., Cayouette, M. & Harris, W.A. In vivo time-lapse imaging of cell divisions during neurogenesis in the developing zebrafish retina. Neuron 37, 597–609 (2003).

54. Nelson, B.R. et al. Genome-wide analysis of Müller glial differentiation reveals a requirement for Notch signaling in postmitotic cells to maintain the glial fate. PLoS One 6, e22817 (2011).

